# Obesity control by SHIP inhibition requires pan-paralog inhibition and an intact eosinophil compartment

**DOI:** 10.1101/2020.09.15.299073

**Authors:** Sandra Fernandes, Neetu Srivastava, Chiara Pedicone, Raki Sudan, Elizabeth A. Luke, Otto M. Dungan, Angela Pacherille, Shea T. Meyer, Shawn Dormann, John D. Chisholm, William G. Kerr

**Author notes:** Contributed equally (John D. Chisholm, William G. Kerr).

## Abstract

Previously a small molecule SHIP inhibitor, K118, was shown to reverse high-fat diet induced obesity and improve blood glucose regulation in obese mice. K118 treatment was also found to increase the frequency and number of IL-4 producing eosinophils in the visceral fat as well two potent immunoregulatory myeloid cell populations: M2-polarized macrophages and myeloid derived suppressor cells (MDSC) suggesting an immune regulatory mechanism. However, the cell(s) or SHIP paralog that should be targeted to improve metabolic regulation was not defined. Here we extend our understanding of how chemical inhibition of SHIP paralogs improves metabolic regulation during excess caloric intake. Here we compare SHIP inhibitors in an obesity prevention model and find that selective inhibitors for SHIP1 or SHIP2 lack the ability to prevent weight gain and body fat accumulation during increased caloric intake. Surprisingly, only pan-SHIP1/2 inhibitors can prevent diet-induced obesity. We confirm that both SHIP1 and SHIP2 must be targeted by showing that dual treatment with the SHIP1 and SHIP2 selective inhibitors can reduced adiposity caused by excess caloric consumption. We also show that pan-SHIP1/2 inhibitors of two different chemical classes can control diet-induced obesity and improve blood glucose regulation. Intriguingly, we find that both classes of pan-SHIP1/2 inhibitors require an intact eosinophil compartment to prevent diet-related fat accumulation demonstrating pan-SHIP1/2 inhibitors act via the VAT innate immune compartment to control adiposity However, improved blood glucose regulation by pan-SHIP1/2 inhibition is not dependent upon eosinophils, indicating a separate mechanism of action for diet-related loss of glucose regulation.

## Introduction

Obesity and its associated metabolic disorders have become a global epidemic affecting millions of people worldwide. It is now appreciated that metabolism and immunity are intertwined as inflammatory macrophages recruited to visceral adipose tissue (VAT) can promote the onset of obesity and insulin resistance.(1–3) In addition, an M1-like macrophage population in VAT has recently been shown to limit the thermogenic capacity of adipocytes through consumption of norepinephrine (NE) produced by sympathetic neurons that innervate VAT.(4–6) A counterbalance to these pro-inflammatory myeloid populations in VAT are immunoregulatory myeloid cells, AAM/M2 macrophages and myeloid-derived suppressor cells (MDSC).(7–13) Treg cells can also oppose inflammatory stressors on adipocytes and thus promote improved control of blood glucose levels, ahough they do not reduce adiposity.(8, 14, 15) Innate lymphocyte type 2 (ILC2) cells present in the VAT can, via production of the TH2 cytokines IL-5 and IL-13, promote increased numbers of eosinophils. IL-4 producing eosinophils are then able to bias myeloid differentiation in the VAT toward immunoregulatory AAM/M2 cells to promote leanness and improve blood glucose control.(16–19) In this mechanism of immune control of obesity M2 macrophages express tyrosine hydroxylase (TH) and produce NE to drive UCP1 expression and thermogenesis by adipocytes.(17) This adrenergic function of M2/AAM cells has recently come under quesiton,(20) although others have independently replicated that M2/AMM cells are capable of induced TH expression at the mRNA and protein level.(21)

The family of SH2-containing inositol phosphatases consists of only two paralogs, SHIP1 (*INPP5D*) and SHIP2 (*INPPL1*). SHIP1 and SHIP2 are currently thought to have largely distinct functions *in vivo* as phenotypes in SHIP1^−/-^ mice are primarily hematologic(22–24) or immunological,(25–30) and consequently life-threatening. Conversely, SHIP2^−/-^ mice are viable and lack demonstrable hematolymphoid abnormalities. Intriguingly, SHIP2^−/-^ mice are resistant to obesity following consumption of a high-fat diet (HFD).(31) However, the cellular and molecular basis for this resistance has not been defined, and could perhaps be immune-mediated as SHIP2 is expressed in both hematopoietic and parenchymal tissues. A small molecule selective inhibitor of SHIP2 has been described and consequently has shown efficacy in murine models of diabetes mice and Alzheimer’s disease where it improved blood glucose control(32) or cognition,(33) respectively. A variety of small molecule inhibitors of SHIP1 and both SHIP1 and SHIP2 (pan-SHIP1/2 inhibitors) have been described as well and also been shown to enable targeting of SHIP1 or both paralogs in vivo with evidence of therapeutic efficacy in pre-clinical models of diseases that including cancer,(34, 35) bone marrow transplantation,(36) blood cell recovery,(37, 38) cancer immunotherapy,(39) lethal infection(40) and reduction of β-amyloid in the CNS.(41) Thus, SHIP1 and SHIP2 have proven to be tractable targets *in vivo* whose selective or dual inhibition has potent physiological effects that can abrogate disease without significant toxicity.(42)

The demonstration that the SHIP inhibitor K118 can reverse obesity following consumption of a HFD,(43) suggested that further analysis of how SHIP inhibition mediates obesity control was merited. In addition, our recent analysis of SHIP inhibitors in tumor immunity showed that compounds with modest selectivity for SHIP1, but which also target SHIP2, like K118, can have distinct activity versus more selective SHIP1 inhibitors like 3AC.(39) In addition, our recent analysis of a broad panel of SHIP inhibitors showed that pan-SHIP1/2 inhibitors was optimal for promoting homeostatic functions of microglia *in vitro* and *in vivo*, including phagocytosis of Aβ42, the primary constituent of amyloid plaques present in Alzheimer’s Disease.(41) Using a similar approach here we dissect whether the activity of SHIP1, SHIP2 or both enzymes should be targeted in order to control obesity and blood glucose in the context of excess calorie intake in wild type mice, but also mice that lack eosinophils. These studies reveal that pan-SHIP1/2 inhibitors are required for effective obesity control and that eosinophils are essential for this antiobesity activity, but not for control of blood glucose levels.

## Results

### The pan-SHIP1/2 inhibitor K118 protects mice from HFD-induced obesity

Previously we found that that K118 was effective at reversing obesity in a treatment setting after mice had already become obese due to consumption of HFD over a 2 month period.(43) We subsequently found that K118 has only modest selectivity for SHIP1 and consequently inhibits both SHIP1 and SHIP2 with significant potency.(39, 41) This led us to question whether solely targeting SHIP1 was required for obesity control. In order to more readily test novel SHIP inhibitors that might protect against diet-induced weight and fat gain, we developed a prevention model where mice are placed on HFD while simultaneously being treated with a candidate SHIP inhibitor (SHIPi) compound. To confirm the validity of this model we introduced mice to consumption of a HFD and initiated K118 treatment in the same week with the same dosing regimen (2X/week, 10mg/kg) as we utilized in the obesity treatment model.(43) We found that K118 treatment in this obesity prevention setting also protected adult mice from significant weight gain(**Fig. 1A**) and, importantly, also prevented increased adiposity (% body fat) despite *ad libitum* consumption of HFD over a 4 week period.(**Fig. 1B**) Importantly we did not observe wasting in the mice due to K118 treatment as their % lean mass was maintained as compared to % lean mass at initiation of HFD consumption.(**Fig. 1C**) As expected, vehicle treated mice gained significant body weight and increased their % body fat and decreased their % lean mass following initiation of HFD consumption.(**Fig. 1A-C**) Consistent with its ability to reverse obesity in a treatment setting,(43) these findings confirmed the pan-SHIP1/2 inhibitor K118 is also able to prevent obesity upon consumption of a HFD and validate the use of this model to assess SHIP inhibition strategies in diet-induced obesity.

**Fig. 1.**
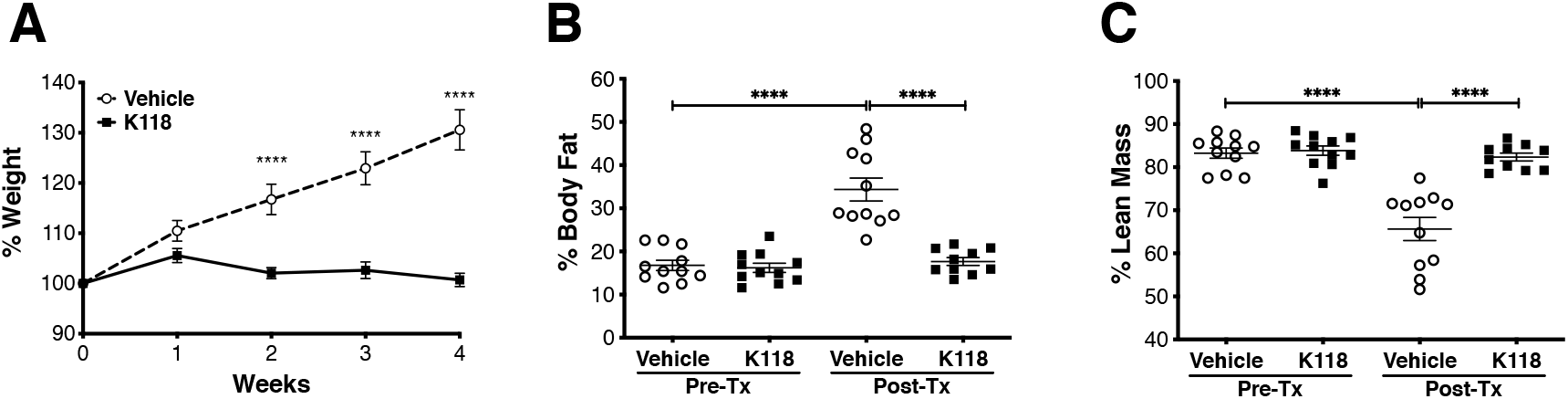
The pan-SHIP1/2 inhibitor K118 prevents the onset of obesity. (A) % Body weight, (B) % body fat and (C) % lean mass measurements on C57BL/6 mice following *ad libitum* consumption of a HFD and simultaneous treatment with K118 or vehicle (H2O). Mice were dosed with K118 two times per week (on days 1 and 4 of each week at 10mg/kg via i.p. injection) for the 4 week duration of the study. Body Fat and Lean mass were measured by DEXA imaging before initiation of the study and after 4 weeks on HFD (Mean±SEM, 2-way Repeated measures ANOVA with Bonferroni multiple comparison test in **A**, two-tailed t-test with Welch’s correction where needed for **B** and **C**, ****p<0.0001, pooled from two experiments with n=5)

### Paralog-selective SHIP inhibitors are ineffective at prevention of diet-induced obesity

The above findings established that a pan-SHIP1/2 inhibitor could prevent obesity following increased caloric intake. However, this could be due to K118 inhibition of either SHIP1 or SHIP2, and thus unrelated to its ability to inhibit both SHIP paralogs simultaneously. If this were the case, then either SHIP1- or SHIP2-selective inhibition should be able to protect mice from diet-induced weight gain and adiposity. In fact, genetic analysis suggested that solely targeting SHIP2 might enable control of diet-induced obesity.(31) To examine these different possibilities, we treated mice placed on a HFD with either a SHIP1-selective inhibitor (3AC)(37) or SHIP2-selective inhibitor (AS194940),(32) both of which have each been shown to be effective at targeting SHIP1 or SHIP2 *in vivo*, respectively.(32, 34, 37–39, 44, 45) We found that both the SHIP1-selective inhibitor 3AC (**Fig. 2A,B,C**) and the SHIP2-selective inhibitor AS1949490 (**Fig. 2D,E,F**) are incapable of preventing body weight gain and increased adiposity (% body fat), and loss of lean mass upon consumption of a HFD. Mice treated with either compound gained essentially the same amount of weight and increased body fat to the same degree as their respective vehicle controls. That SHIP paralog selective inhibitors fail to protect from obesity suggested that the capacity of K118 to prevent obesity could be due to its capacity inhibit both SHIP1 and SHIP2 simultaneously.

**Fig. 2.**
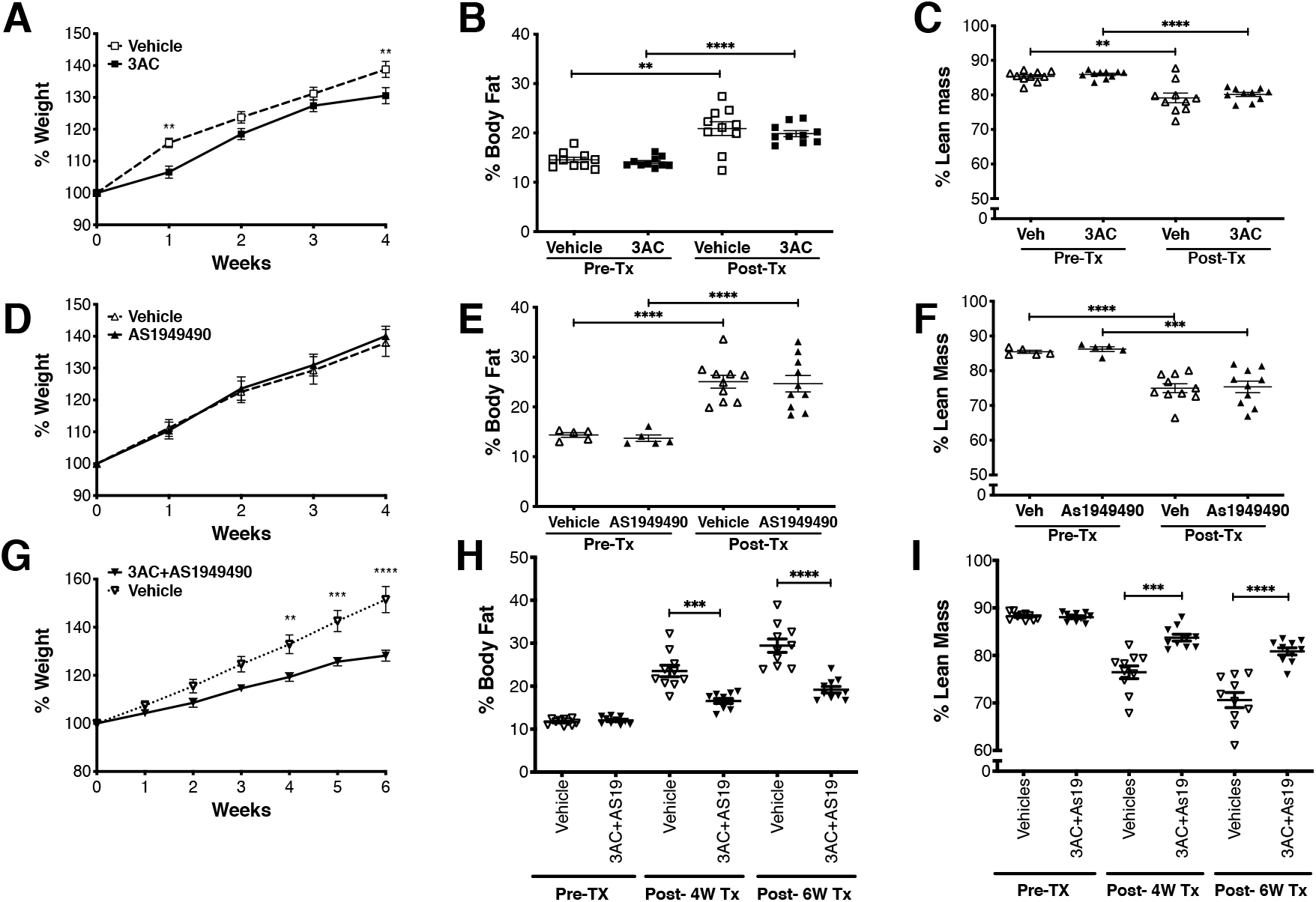
SHIP paralog-selective inhibitors are unable to prevent obesity, but dual treatment with SHIP1- and SHIP2-selective inhibitors reduces weight and body fat increases due to increased caloric intake. **(A, D, G)** % Body weight, **(B, E, H)** % body fat and (**C, F, I**) % lean mass in C57BL/6 mice during HFD consumption and simultaneous treatment with (**A-C**) 3AC or vehicle (0.3% Klucel:saline), (**D-F**) AS1949490 or vehicle (5% DMSO:saline) or (**G-I**) dual treatment with both 3AC and As1949490 (As19) or dual vehicle (0.3% Klucel:saline + 5% DMSO:saline). Mice were dosed with 3AC and/or AS1949490 or vehicle two times per week (on days 1 and 4 of each week at 26.5mg/kg for 3AC or 20mg/kg for As1949490 via i.p. injection) for the 4 week duration of the study. (Body Fat and Lean mass were measured by DEXA imaging before initiation of the study and after 4 or 6 weeks on HFD with SHIPi or vehicle treatment (Mean±SEM, 2-way Repeated measures ANOVA with Bonferroni multiple comparison test in **A, D, G**, two-tailed t-test with Welch’s correction when needed for **B, C, E, F, H,** and **I**, **p<0.01, ***p<0.001, ****p<0.0001, pooled from two experiments with n=5).

### Dual targeting of SHIP1 and SHIP2 with paralog-selective inhibitors reduces obesity and loss of lean mass caused by increased caloric intake

We considered then that both SHIP paralogs must be targeted *in vivo* in order to reduce weight gain and increased adiposity during HFD consumption. Thus, we undertook a study where we simultaneously treated mice consuming a HFD with both 3AC and AS1949490. In this setting we found that both vehicle and 3AC+AS1949490 treated groups gained weight while consuming a HFD. However, the 3AC+AS1949490 treated group gained significantly less weight at 4 weeks after initiation of HFD. When inhibitor treatment was continued through 6 weeks the difference in body weight became even more apparent.(**Fig. 2G**) Importantly, mice co-treated with 3AC+AS1949490 also acquired considerably less body fat than vehicle treated mice as measured at 4 and 6 weeks of HFD consumption,(**Fig. 2H**) and retained a greater proportion of lean mass vs. the vehicle control group.(**Fig. 2I**) As with body weight, the reduction in body fat was more apparent after the treatment was continued for 6 weeks. While 3AC+AS1949490 treatment does provide a significant degree of protection from diet-induced obesity this protection is not as complete as we observed with K118 (**Fig. 1**). The reason for this is not clear, but one possibility is that K118 has greater potency against SHIP1 than 3AC, and thus more effectively targets both SHIP paralogs than the combination of 3AC and AS1949490. Taken together these findings demonstrate that significant control of diet-induced obesity is achieved by simultaneous inhibition of both SHIP paralogs, and not by paralog selective SHIP inhibition.

### Other pan-SHIP1/2 inhibitory compounds also reduce diet-induced obesity

To further test and validate the hypothesis that pan-SHIP1/2 inhibition is necessary for effective obesity control we sought to test other pan-SHIP1/2 inhibitors we have recently identified.(41, 42) These include compounds of a completely different chemical class than aminosteroids such as the tryptamine K149,(34, 39, 46) but also a water soluble aminosteroid K161 that has significant bioavailability *in vivo*, including the ability to cross the blood brain barrier to access brain-resident microglia.(41) To provide confirmation of the utility of pan-SHIP1/2 inhibitors as effective therapeutics for obesity control we examined whether K149 could also prevent or reverse obesity caused by consumption of a HFD.(**Fig. 3**) As with the aminosteroid K118, the tryptamine K149 significantly reduced weight gain during a 6 week period of HFD consumption with mice only gaining a small amount of weight.(**Fig. 3A**) This was reflected in the K149-treated mice exhibiting only a negligible, but significant, increase in % body fat, while vehicle treated mice gained significantly more body fat than their K149-treated counterparts.(**Fig. 3B**) Importantly, K149 treated mice on HFD maintained the same % lean mass as they had prior to starting on HFD,(**Fig. 3C**) indicating K149 does not promote weight control through adverse effects on physiology that promote wasting. We also found that K149 can protect mice in a treatment model of HFD induced obesity. In this setting, mice were first rendered obese by consumption of HFD for 2 months after which they were treated with K149 or vehicle (2X per week) whilst continuing to consume a HFD. We found that K149 significantly decreased both body weight (**Fig. 3D**) and percent body fat (**Fig. 3E**) within 4 weeks of treatment despite continued HFD consumption. Importantly, K149 treatment increased % lean mass (**Fig. 3F**) in the obese mice indicating weight loss was not due to wasting. We also found that the pan-SHIP1/2 inhibitory aminosteroid K161(41) also protects mice from weight gain and obesity in a prevention setting,(**Fig. 4A-C**) but also when used to treat mice that are already obese and continue to consume a HFD diet.(**Fig. 4D-F**) Thus, three pan-SHIP1/2 inhibitory compounds (K118, K149, K161) derived from two different chemical classes (aminosteroids and tryptamines) have a potent capacity to mediate obesity control in the context of excess caloric intake, while SHIP paralog selective inhibitors like 3AC and AS1949490 are unable to protect from diet-induced obesity.

**Fig. 3.**
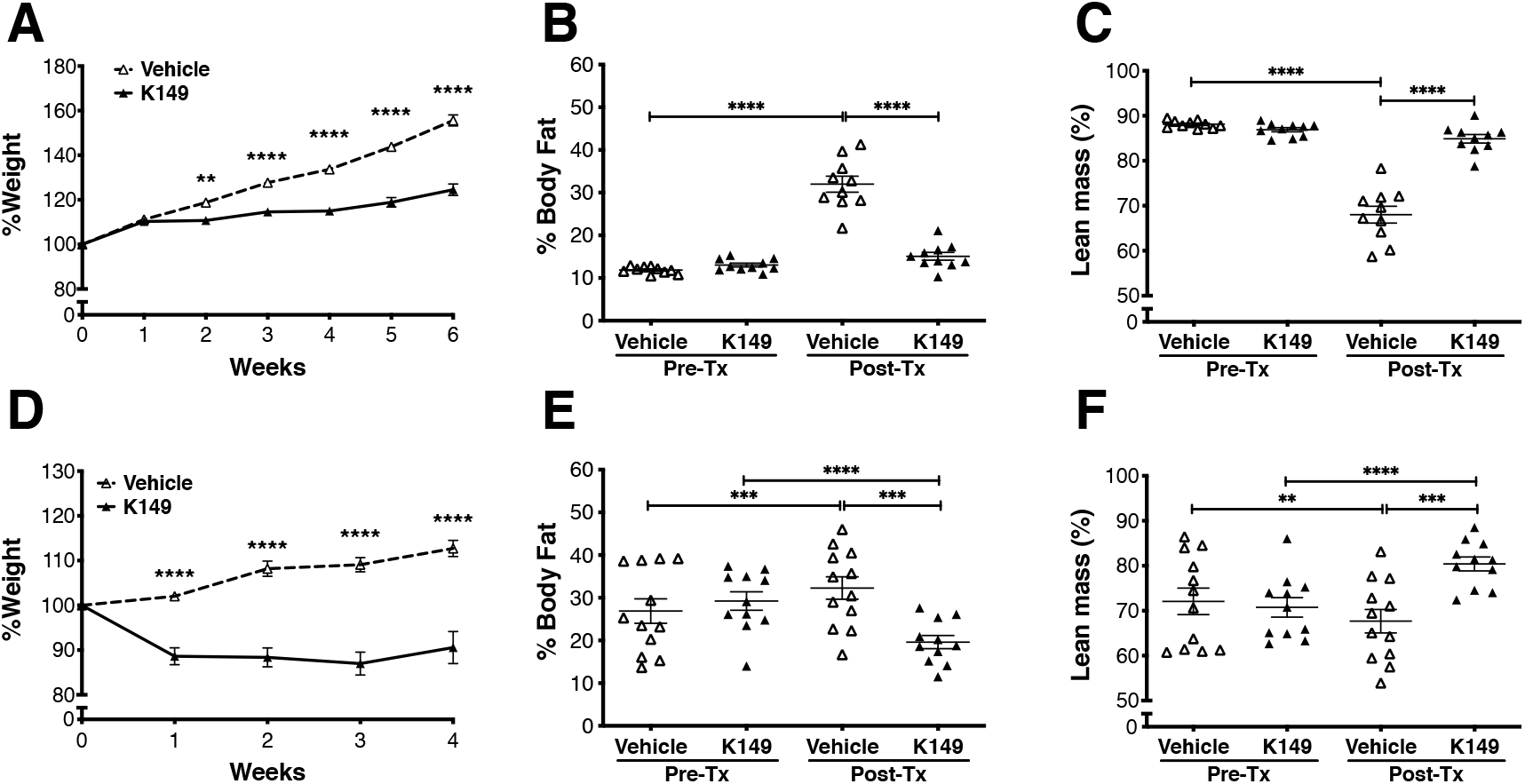
The pan-SHIP1/2 inhibitor K149 prevents the onset and reverses diet-induced obesity. (**A, D**) % Body weight, (**B, E**) % body fat and (**C, F**) % lean mass measurements on C57BL/6 mice following *ad libitum* consumption of a HFD and simultaneous treatment with K149 or vehicle (5%DMSO:saline). Mice were dosed with K149 two times per week (on days 1 and 4 of each week at 10mg/kg via i.p. injection) for the 4-6 week duration of the study. Mice in **A-C** were placed on HFD at the time of the first dose of SHIPi or vehicle, while mice in **D-F** were placed on HFD for 8 weeks prior to the start of SHIPi or vehicle and maintained on the diet for the 4 weeks of the study. Body Fat and Lean mass were measured by DEXA imaging before initiation of the treatment and after 4 and 6 weeks on HFD (Mean±SEM, 2-way Repeated measures ANOVA with Bonferroni multiple comparison test in **A, D** two-tailed t-test for **B, C, E** and **F**, **p<0.01, ***p<0.001,****p<0.0001, each model (prevention or reversal) is pooled from two experiments with n=5).

**Fig. 4.**
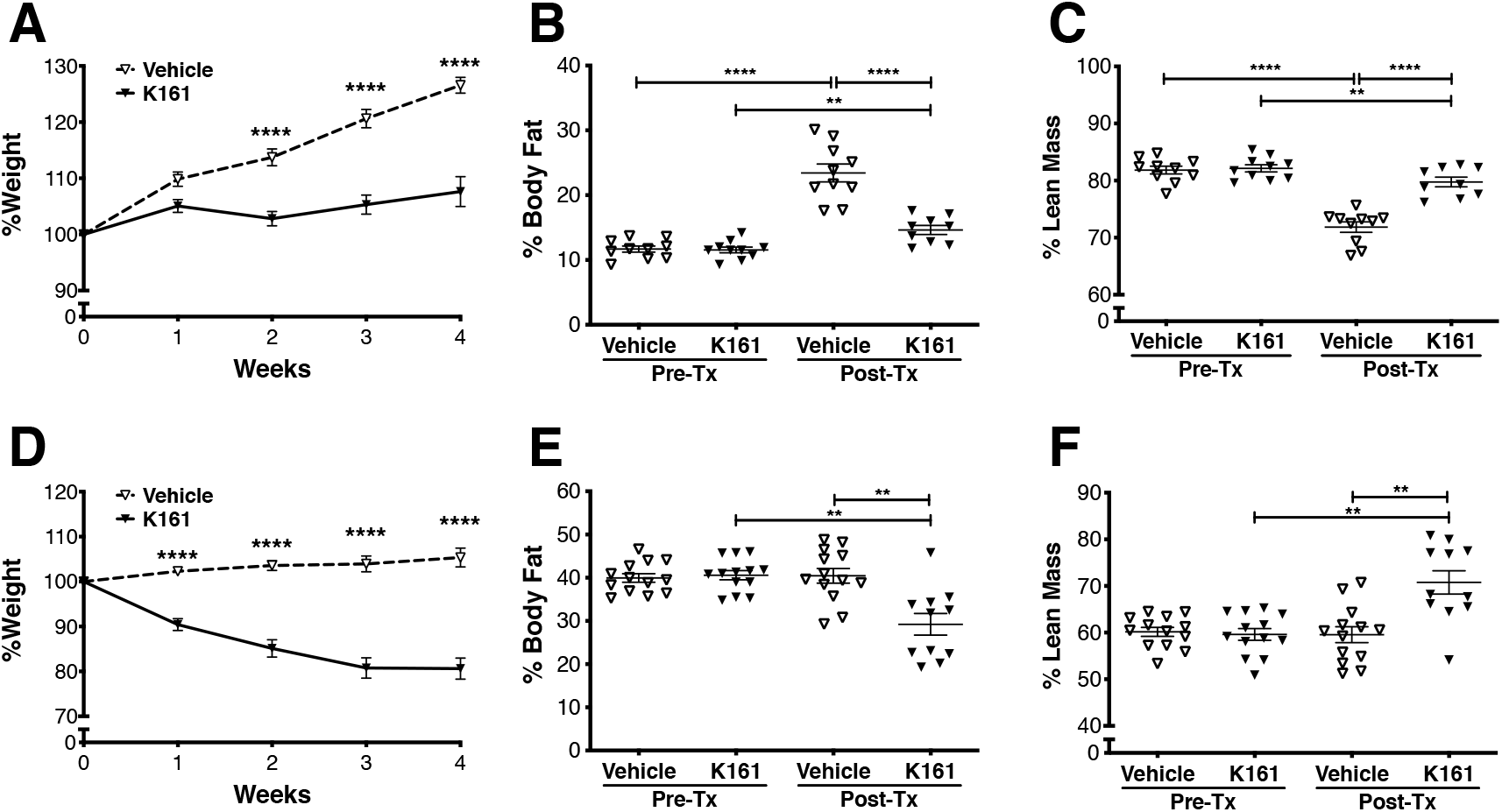
The pan-SHIP1/2 inhibitor K161 prevents the onset and reverses Diet-induce obesity. (**A, D**) % Body weight, (**B, E**) % body fat and (**C, F**) % lean mass measurements on C57BL/6 mice following *ad libitum* consumption of a HFD and simultaneous treatment with K161 or vehicle (H2O). Mice were dosed with K161 two times per week (on days 1 and 4 of each week at 10mg/kg via i.p. injection) for the 4 week duration of the studies. Mice in **A-C** were placed on HFD at the time of the first dose of SHIPi or vehicle, while mice in **D-F** were placed on HFD for 8 weeks prior to the start of SHIPi or vehicle and maintained on the diet for the 4 weeks of the study. Body Fat and Lean mass were measured by DEXA imaging before initiation of the treatment and after 4 weeks on HFD (Mean±SEM, 2-way Repeated measures ANOVA with Bonferroni multiple comparison test in **A, D**, two-tailed t-test for **B, C, D** and **F**, **p<0.01,****p<0.0001, each model (prevention or reversal) is pooled from two experiments with n=5).

### Other pan-SHIP1/2 inhibitory compounds promote increased eosinophil function in the face of HFD stress

The pan-SHIP1/2 inhibitor K118 was found to reduce SHIP1 expression in VAT-resident eosinophils and increase the frequency of IL-4-producing eosinophils in the VAT of obese mice.(43) Thus, we assessed whether the novel pan-SHIP1/2 inhibitors that mediate obesity control or reversal, K149 and K161, can protect the eosinophil compartment and its’ IL-4 producing function during the adipogenic stress associated with excess caloric intake. In the setting of obesity prevention K149 increases the frequency of eosinophils producing IL-4 vs. vehicle controls following 6 weeks of sustained HFD consumption (**Fig. 5A**) and surprisingly increased the average IL-4 made by these cells based on intracellular staining for IL-4.(**Fig. 5B**) Increased IL-4 production by the eosinophil compartment was found to correlate with an M2/AAM polarized macrophage compartment and increased MDSC in the VAT of the K149 treated mice,(**Fig. 5C**) consistent with the effects of IL-4 on VAT macrophages.(11, 16, 47) We assessed the same immune components in mice rendered obese by HFD consumption mice, but treated with the water soluble aminosteroid pan-SHIP1/2 inhibitor K161. K61 treatment demonstrated a clear capacity to promote VAT eosinophil function in the face of adipogenic stress as K161 increased both the frequency of total eosinophils and the IL-4 producing eosinophil subset,(**Fig. 5D**) as well as their production of IL-4 on a per cell basis in the obesity treatment model(**Fig. 5E**). As with K118 and K149, K161 also polarizes the VAT macrophage compartment toward M2 macrophages and increases MDSC numbers in the VAT.(**Fig. 5F**) Thus, pan-SHIP1/2 inhibition has the unique capacity to promote immune features in the VAT that are correlated with maintenance of the lean state.

**Fig. 5.**
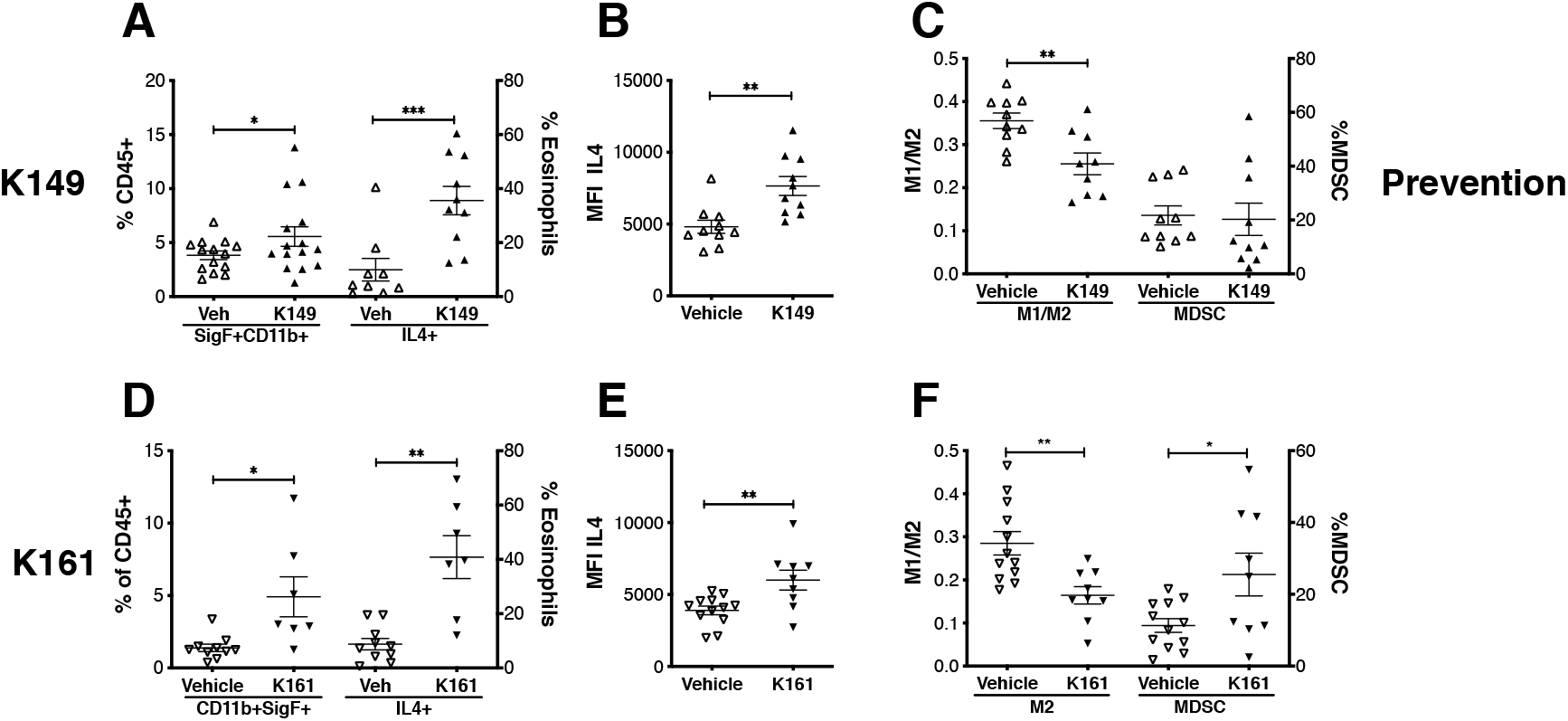
SHIPi leads to increase in the frequency of IL4+ and total IL4 in Eosinophils, promotes M2/AAM polarization and increases MDSC in eWAT. (**A, D**) Frequency of IL4+ expressing and (**B, E**) total intracellular IL4 was increased Eosinophils (CD45+SigF+CD11b+) isolated from eWAT after treatment with pan-SHIP1/2 inhibitors. (**C, F**) Polarization of macrophages to an M2/AAM phenotype (see representative electronic gating used to estimate M1 (CD11b^+^F4/80^+^CD86^lo^CD11c^+^) or M2/AAM cells (CD11 b^+^F4/80^+^CD86^lo^CD11c^−^) and MDSC (CD11+Gr1+) were increased was observed in eWAT after SHIPi treatment. (**A-C**, prevention model: K149 treatment was initiated on the same day the mice were place on HFD and maintained for the 6 weeks of the study. **D-F** obesity reversal model, mice were place on HFD for 8 weeks prior to initiation of the 4-weel K161 treatment. Mean±SEM, one-tailed t-test, *p<0.05, **p<0.01, ***p<0.001 each model (prevention with K149 or reversal with K161) is pooled from at least two experiments with n=5). *Please see Supplemental Figure 1A-D for representative examples of flow cytometric electronic gates used to estimate the frequency of total VAT eosinophils, IL4-producing eosinophils, IL4 MFI in eosinophils, M1/M2 polarization and MDSC frequencies*.

### Pan-SHIP1/2 inhibitors are unable to reduce obesity and polarize the macrophage compartment in mice that lack an intact eosinophil compartment

Mice homozygous for the ΔdblGATA mutation lack eosinophils due to a requirement by hematopoietic stem cells for eosinophil lineage commitment promoted by the GATA1 transcription factor.(48) ΔdblGATA mice were subsequently used to show that an eosinophil compartment is required for protection from weight gain and loss of glucose homeostasis following consumption of a HFD.(16, 49) Because multiple pan-SHIP1/2 inhibitors promote VAT eosinophil function in different diet-induced obesity settings, we hypothesized that eosinophils might be a critical cellular target of pan-SHIP1/2 inhibitors for obesity control. Thus, we tested the capacity of K118 to mediate obesity control in ΔdblGATA mice and BALB/C mice of the same genetic background. When BALB/C mice, which have an intact eosinophil compartment, were placed on HFD and simultaneously treated with K118 they did not gain body weight to the same degree as vehicle treated controls.(**Fig. 6A**) They also showed no increase in % body fat (**Fig. 6B**) and their lean mass was comparable to that prior to the initiation of HFD 6 weeks earlier. As expected, vehicle treated BALB/C mice showed a dramatic increase in % body fat and loss of lean mass after 6 week of HFD consumption.(**Fig. 6C**) In the ΔdblGATA mice placed on a HFD K118 treatment surprisingly did reduce the amount of body weight relative to vehicle treated ΔdblGATA mice placed on a HFD.(**Fig. 6A**) However, the K118 treated ΔdblGATA mice increased the % body fat to the same degree as vehicle treated ΔdblGATA mice,(**Fig. 6B**) indicating eosinophils are required for K118 to prevent diet-induced increases in adiposity during excess caloric intake. Intriguingly, K118 treated ΔdblGATA mice retained a comparable % lean mass while their vehicle treated counterparts lost significant % of lean body mass.(**Fig. 6C**) To provide an independent measure for loss of eosinophil function, as further evidence that K118 acts on eosinophils *in vivo*, we also assessed M2 macrophage polarization in the vehicle and K118 treated ΔdblGATA mice on HFD. We found that that M2/AMM polarization was not induced in ΔdblGATA mice by K118 as compared to vehicle controls.(**Fig. 6D**) The frequency of MDSC was still increased indicating the effect of K118 on MDSC numbers in the VAT is independent of its effect on eosinophils. These results also indicate increased MDSC numbers in the VAT associated with pan-SHIP1/2 inhibition are not required for obesity control.(**Fig. 6D**) We then performed a similar study with both K149 and K161. Mice that consumed a HFD for 6 weeks while being treated with K149 (**Fig. 6E**) or K161(**Fig. 6I**) lost weight as compared to vehicle controls, but did not show a significant increase in % body fat (**Fig. 6F, J**) or a decrease in % lean mass as compared to their respective vehicle control groups following 6 weeks of HFD consumption. In addition, both K149 (**Fig. 6H**) and K161 (**Fig. 6L**) treatment of mice consuming HFD for 6 weeks failed to polarize the VAT M1/M2 compartment toward M2 cells, but did show significantly increased frequency of MDSC present in the VAT. Thus, all three pan-SHIP1/2 inhibitors tested, K118, K149, K161, fail to control obesity and polarize the macrophage compartment toward M2 cells in mice that lack an eosinophil compartment during consumption of a HFD.

**Fig. 6.**
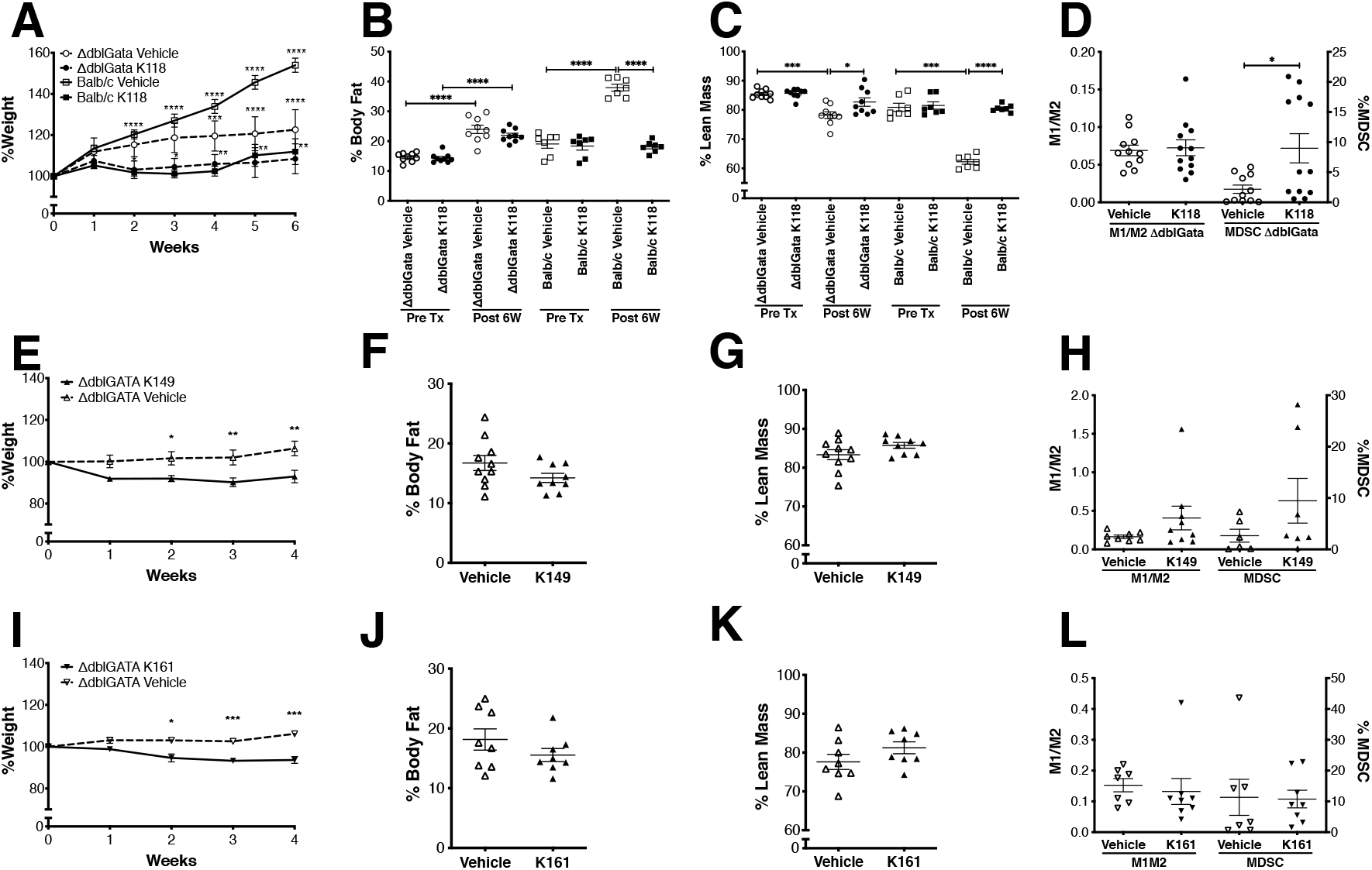
Pan-SHIP1/2 inhibitors limit obesity and promotes M2/AAM polarization via the eosinophil compartment. **(A)** ΔdblGATA and Balb/c mice were placed on HFD for 6 weeks and simultaneously treated with K118 or vehicle twice a week as described above. Body weight was measured on a weekly basis after initiation of HFD consumption. **(B)** The % body fat and (**C**) % lean mass was determined by DEXA imaging of the K118 or Vehicle treatment groups in (A) before (Pre Tx) and after (Post 6W) 6 weeks of HFD consumption. **(D)** eWat macrophages are not polarized to an M2/AAM phenotype (from mice in **A-C**) in ΔdblGATA following simultaneous HFD and 6 week treatment of K118 as determined by flow cytometry of eWAT SVC fraction as in Figure 5. (**E-L**)ΔdblGATA mice were placed on HFD for 8 weeks prior to the start of SHIPi or vehicle and maintained on the diet for the 4 weeks of the study. (**E, I**) % Body weight, (**F, J**) % body fat and (**G, K**) % lean mass measurements on ΔdblGATA treated two times per week as above with (**E-H**) K149 or vehicle (5%DMSO:saline) or (**I-L**) K161 or vehicle (H2O). Body Fat and Lean mass were measured by DEXA imaging before initiation of the SHIPi treatment and after 4 weeks on HFD. (**H, L**) eWat macrophages are not polarized to an M2/AAM phenotype in ΔdblGATA mice following K149 or K161 treatment as determined by flow cytometry of eWAT SVC fraction as above. (Mean±SEM, 2-way Repeated measures ANOVA with Bonferroni multiple comparison test in **A, E, I**, two-tailed t-test for **B-D, F-H, J-L** *p<0.05, **p<0.01, ***p<0.001, ****p<0.0001, each model is pooled from two experiments).

### Pan-SHIP1/2 inhibitors mediate control of blood sugar levels independent of their effect on adiposity and eosinophils

In our previous study we found that K118 also improves blood glucose regulation in obese mice.(43) We found this also to be the case with the pan-SHIP1/2 inhibitors K149(**Fig. 7A**) and K161(**Fig. 7C**) as they both reduced blood glucose levels in mice after 6 weeks of HFD consumption as compared to vehicle controls. We also tested this in the setting of obesity treatment with K161, where mice become obese following 2 months of HFD consumption and then are treated for two weeks with K161 or vehicle while continuing to consume HFD. We found that K161 also mediated a significant reduction in blood glucose levels vs. vehicle controls.(**Fig. 7E**) We speculated this might have simply been a byproduct of obesity control. However, this appears to be an entirely independent effect of pan-SHIP1/2 inhibition as K149 (**Fig. 7B**) or K161(**Fig. 7D**) treated ΔdblGATA mice still show significant reductions in blood glucose vs. vehicle controls in an obesity prevention setting, despite the fact that % body fat is not decreased in ΔdblGATA vs vehicle controls treated with either K149 or K161.(**Fig. 6**) We also found that K161 significantly reduced blood glucose levels in obese ΔdblGATA mice. (**Fig. 7F**) Thus, pan-SHIP1/2 inhibitors can improve blood glucose control in the setting of increased caloric intake and frank obesity, and do so independent of their ability to reduce adiposity via the eosinophil compartment.

**Fig. 7.**
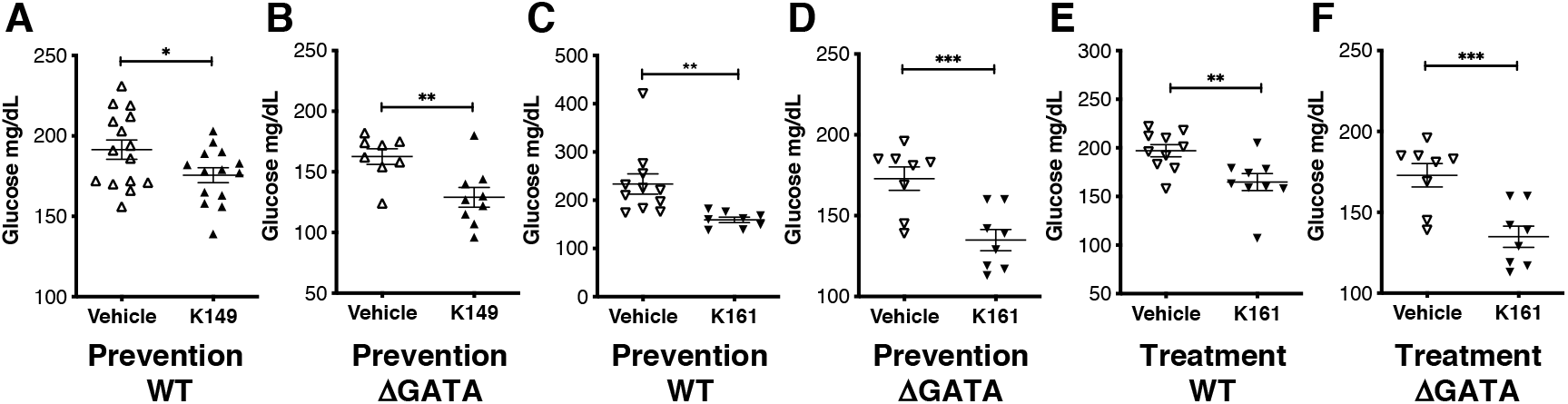
Pan-SHIP1/2 inhibitors mediate control of blood sugar levels independent of their effect on adiposity and eosinophils. End-point *ad libitum* blood glucose levels were measured in (**A, C, E**) C57BL/6 or (**B, D, F**) ΔdblGATA mice following treatment with pan-SHIP1/2 inhibitors (**A, B**) K149 or vehicle (5%DMSO:saline) or (**C-F**) K161 or vehicle (H2O). Measurements we performed 16h after the final dose of SHIPi from mice treated above. SHIPi was initiated on the same day that mice were place on HFD for the prevention model (**A-D**) while mice were place on HFD for 8 weeks prior to initiation and maintainded for the duration SHIPi in treatment model (**E, F**).

## Discussion

Here we show that SHIP inhibitory compounds capable of inhibiting both SHIP1 and SHIP2 are optimal for reducing obesity associated with excess caloric intake. We show that this is the case for both aminosteroid- and tryptamine-based pan-SHIP1/2 inhibitors. However, compounds that only selectively inhibit SHIP1 or SHIP2 are completely ineffective at preventing accumulation of excess body fat associated with excess caloric intake and yet when these same two paralog-selective inhibitors are coadministered to mice consumption of a HFD the combination protects from increased accumulation of body fat. We find that an eosinophil compartment is required for promotion of the lean state by pan-SHIP1/2 inhibitors. This activity is strongly correlated with the ability of all pan-SHIP1/2 inhibitors to increase both the frequency of IL4-producting eosinophils, the amount of IL4 they produce per cell and polarization of the VAT macrophage compartment toward M2 macrophages. Surprisingly, pan-SHIP1/2 inhibitors ability to improve blood glucose control is not linked to their capacity to prevent increased body fat accumulation, indicating their ability to improve glucose control has a distinct mechanism of action.

Control of both obesity and blood glucose levels by immune cells present in the VAT is now understood to have a prominent role in metabolic control, and also its dysregulation.(1–3, 7–19, 21) The recently described M1-like inflammatory macrophage subsets in VAT that can act as norepinephrine (NE) sinks may have provided the much sought after ‘missing-link’ between VAT immune function and energy utilization by lipid-laden adipocytes.(4–6) However, direct production of NE by M2 macrophages to stimulate lipolysis and thermogenesis by VAT adipocytes remains a distinct possibility.(17, 21) In fact, there is likely a competition between these distinct macrophage functions that ultimately types the balance toward metabolic control or dysregulation sustained excess caloric intake. Of the cells present in the VAT immune milieu, we find that pan-SHIP1/2 inhibition requires an eosinophil compartment to mediate obesity control. How pan-SHIP1/2 inhibitors achieve this effect on adiposity via eosinophils remains to be defined. However, pan-SHIP1/2 inhibition was consistently found to increase the frequency of IL4-producing eosinophils in the VAT as well as the amount of IL4 they produce on a per cell basis. IL4 production by VAT eosinophils can drive polarization of the local myeloid compartment in VAT toward cells of an M2 phenotype(16) and M2 polarization was consistently observed with all three pan-SHIP1/2 inhibitors tested in HFD obesity models used here. Whether this also leads to catecholamine synthesis by M2 cells remains to be determined. Another possibility, however, is that M2 polarization depletes the inflammatory M1 macrophages in VAT that can sequester NE from adipocytes and thereby increases β-adrenergic signaling in the adipocyte compartment indirectly. One or both are possibilities might be explored to better understand how pan-SHIP1/2 inhibition mediates obesity control via the eosinophil compartment and their influence on macrophage differentiation in the VAT.

We had previously shown that K118 improves blood glucose control, including the ability of the host to sequester glucose from the plasma.(43) We found that the more potent and water-soluble aminosteroid pan-SHIP1/2 inhibitor K161 and the tryptamine K149 also improved blood glucose control in both HFD obesity models. However, it was surprising to find pan-SHIP1/2 inhibitors mediate blood glucose control in ΔdblGATA mutant mice that lack eosinophils which are resistant to obesity control by pan-SHIP1/2 inhibitors. This indicates a separate and distinct mechanism of action for blood glucose control by pan-SHIP1/2 inhibitors that does not require eosinophil function and their polarization of the macrophage compartment toward an M2 phenotype. Toward that end, it is possible that blood glucose control by pan-SHIP1/2 inhibitors could be acting via the VAT Treg compartment. The VAT Treg compartment exerts a profound control over blood glucose levels in obesity.(8, 15) In our previous study, we found that the pan-SHIP1/2 inhibitor K118 prevents progressive diminution of tissue-resident Treg cells in the VAT during obesity.(43) Thus, it is possible that these compounds can act to prevent the demise of tissue-resident Treg compartment in the VAT of obese mice to maintain or improve blood glucose regulation. Consistent with this hypothesis, genetic SHIP1 deficiency increases the number and function of Treg cells in the spleen and lymph nodes through both cell-intrinsic and −extrinsic pathways.(30, 50) Alternatively, this could be due to the SHIP2 inhibitory effect of pan-SHIP1/2 inhibitors as the SHIP2 selective inhibitor AS19490 improves glucose control in diabetic *db/db* mice(32) and SHIP2^−/-^ mutant mice maintain normal glucose control when placed on a HFD.(31)

Obesity and the metabolic dysregulation that accompanies it are major emerging challenges for global health. Thus, novel treatments are needed to combat obesity and its diabetic consequences. We have shown here that multiple SHIP1/2 inhibitors of two different chemical classes not only combat body fat accumulation associated with excess caloric intake, but also improve the control of plasma glucose levels. Intriguingly, they appear to act primarily by promoting homeostatic function of innate immune cells found in VAT, although their impact on glucose control may be due effects on non-immune cells that express only the SHIP2 paralog. Further research is thus needed to increase our understanding of how these compounds achieve both of these metabolic effects. Nonetheless, pan-SHIP1/2 inhibitory small molecules represent promising candidates for drug development in both obesity and diet-induced diabetes.

## MATERIALS & METHODS

### Mice

C57BL6/J (Stock No.:000664), Balb/c (Stock No.:000651) and ΔdblGATA (Stock No.: 005653) mice were purchased from Jackson Labs. Mice were fed a high fat 60 kcal% diet (HFD, Research Diets D12492) *ad libitum* starting at 6 weeks of age and were maintained on this diet for the duration of the study. For the prevention studies, mice were treated with SHIPi starting on day 1 of HFD. For the obesity reversal studies, mice were placed on HFD for 8 weeks prior to initiating the SHIPi treatment. The SUNY Upstate Medical University IACUC approved all animal experiments.

### SHIP inhibitor treatment of mice

K118 and K161 were dissolved in water and injected at 10mg/kg. AS1949490 (Tocris Bioscience, Minneapolis, MN) and K149 were initially dissolved in DMSO and diluted to 5% with normal saline (0.9%) and administered at 20mg/kg and 10mg/kg, respectively. 3AC was emulsified in 0.3% Klucel (hydroxypropylcellulose M.W. 60,000, Sigma Aldrich, USA): saline (w/v) and administered at 26.5mg/kg. The vehicle groups were H2O (10ul/g) for K118 and K161, 5% DMSO: saline (10ul/g) for As1949490 and K149, or 0.3% Klucel:saline (20ul/g) for 3AC. Mice were injected i.p. with the drug or vehicle twice a week for two to six weeks as indicated in the respective figures.

### Isolation of Immune cells from adipose tissue and flow cytometry

Perigonadal white adipose tissue (WAT) was digested in 6 ml of Collagenase II buffer (Collagenase II 2mg/ml; Sigma Aldrich, USA, 0.5 % BSA in 1X PBS) at 37 °C with shaking at 200 r.p.m for 20 min. Digested tissue was filtered through 100μm filter and centrifuged at 500g for 10 min. Floating adipocytes were removed and the pellet containing stromal vascular fraction (SVF) was resuspended in 2ml of RBC lysis buffer (Thermo Fisher Scientific). Cells were washed and Fc blocked (TruStain FcX(anti-mouse CD16/32) antibody, Biolegend, San Diego, CA) followed by standard flow cytometry surface staining using fluorochrome conjugated antibodies for 30 min on ice. Cells were washed twice and stained with Zombie Aqua Live/Dead (Biolegend, San Diego, CA) stain kit for 20 min on ice. For intracellular staining, cells were fixed with IC Fixation Buffer (Thermo Fisher Scientific) for 20 min on ice and washed with 1X Permeabilization buffer (Thermo Fisher Scientific). Cells were Fc blocked as above in 1X Permeabilization buffer and stained in 1X Permeabilization buffer for intracellular antigens on ice for 30 mins, and washed twice in 1X Permeabilization buffer. Following combination of antibodies were used for staining, anti-mouse CD19-BV711, CD11b-APC-Cy7, F4/80-AlexaFluor488, Gr1-BV605, CD11c-PE-Cy7, CD86-BV786, CD45-BV786, IL4-AlexaFluor488 (Biolegend, San Diego, CA), Gr1-APC-Cy7, Gr1-PerCP-Cy5.5, SigF-AlexaFluor647, SigF-PE, CD45-PerCP-Cy5.5 (BD Biosciences, San Jose, CA). Stained cells were acquired on a BD Fortessa flow cytometer (BD Biosciences, San Jose, CA) and data were analyzed using FlowJo software version 9.9.6 and 10.6 (BD Biosciences, San Jose, CA).

### Blood glucose

Blood glucose was measured in whole blood from tail vein using a blood glucometer (Accu-Chek, Roche Diagnostics, Indianapolis, IN), 16 hours after the final dose of SHIPi in each study.

### DEXA analysis

Body composition was measured pre and post treatment with a PIXImus2 densitometer (GE, Madison, WI), and analyzed with software version 2.10. Mice were anesthetized with isoflurane (Forane, USP, Baxter Healthcare Corporation, IL) throughout body scan.

### Statistical analysis

All statistical analyses were performed using the statistical software Prism 8.4 (GraphPad, San Diego, CA) and is presented as mean ± SEM. Student t-test were used for single variables or two-ANOVA followed by Bonferroni post-test for multiple variables, as described in the legend to each figure.

## Author Contributions

S.F., N.S., C.P., J.D.C. and W.G.K designed, analyzed, wrote and edited the manuscript. S.F., N.S., C.P. R.S., and E.L. performed all obesity related experiments, analyzed data and generated figures. O.M.D., A.P., S.T.M. and J.D.C. synthesized 3AC, K118, K161 and K149 used in these studies.

## Acknowledgements

This work was previously supported in part by the Paige Arnold Butterfly Run. WGK was the Murphy Family Professor of Children’s Oncology Research and an Empire Scholar of the State University of NY during early phase of this study. Additionally, W.G.K, S.F. and J.D.C. receive support from NIH grants RO1 AG059717 and R41 HL142451-01A1.

## Notes

**Conflict of interest statement**, W.G.K, S.F., C.P. and J.D.C have patents, pending and issued, concerning the analysis and targeting of SHIP1 and SHIP2 in disease. W.G.K. is Chief Scientific Officer and J.D.C. serves on the Scientific Advisory Board of Alterna Therapeutics which is devoted to developing and commercializing SHIP inhibitor therapeutics and holds equity. The other authors have no conflicts to disclose.

### Competing Interest Statement

W.G.K, S.F., C.P. and J.D.C have patents, pending and issued, concerning the analysis and targeting of SHIP1 and SHIP2 in disease. W.G.K. is Chief Scientific Officer and J.D.C. serves on the Scientific Advisory Board of Alterna Therapeutics which is devoted to developing and commercializing SHIP inhibitor therapeutics and holds equity. The other authors have no conflicts to disclose.

