## Supplemental Figure 1A_D for "Obesity control by SHIP inhibition requires pan-paralog inhibition and an intact eosinophil compartment"

### Supplemental Figure Legend

**Supplemental Fig. 1. Gating strategy for flow cytometry analysis of SVC from eWAT from C57Bl/6 mice treated with SHIPi.** Doublets and dead cells (stained with Zombie Aqua) were excluded from all analysis. (**A, C.)** Eosinophils were defined as $CD45^+SSC^{hi}$, $CD11b^{lo}$, Siglec-$F^+$, with additional subgate on $IL4^+$ cells (from intracellular staining). Fluorescence minus one (FMO) for IL4 is also shown and was used as a reference point for $IL4^+$ gate placement. (**B, D**) MDSC (left panels) were defined as Live $CD11b^+Gr1^+$. Live cells were then gated for $CD19^-CD11b^+$, then $F4/80^+Gr1^-$. M1 were defined in the subgate as $CD11c^+CD86^{lo}$ and M2 as $CD11c^-CD86^+$ (right panels). M1/M2 was calculated as the frequency of M1 over M2 from parent gate. In the prevention model shown for K149 (**A, B**), mice were place on HFD on the first day of SHIPi treatment and maintained on HFD for 6 weeks. In the treatment model shown for K161(**C, D**) mice were place on HFD 8 weeks prior to the start of SHIPi treatment and maintained on HFD for the 4 week duration of the study.

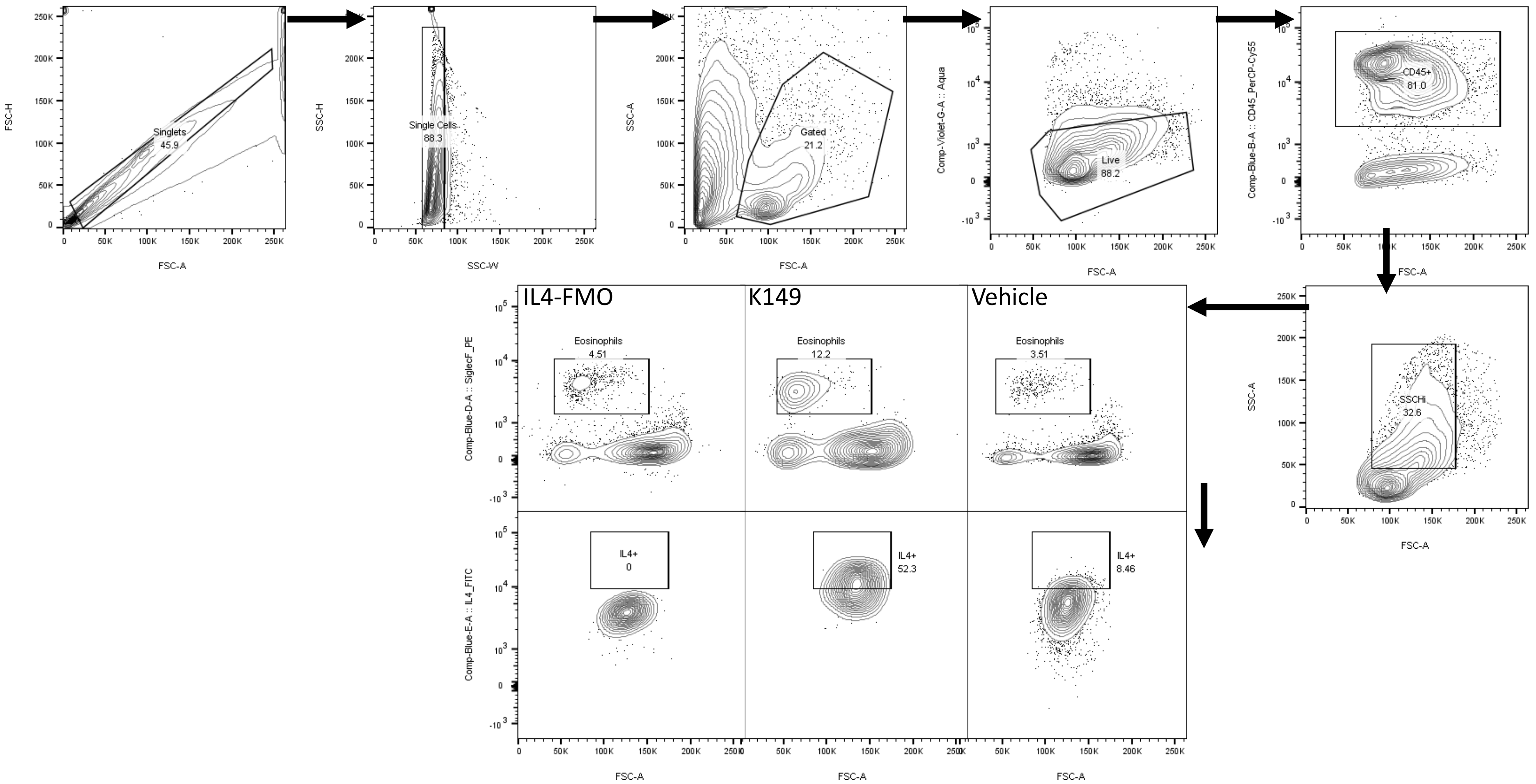


Supplemental Fig.1A Gating strategy Eosinophils K149 Prevention Model

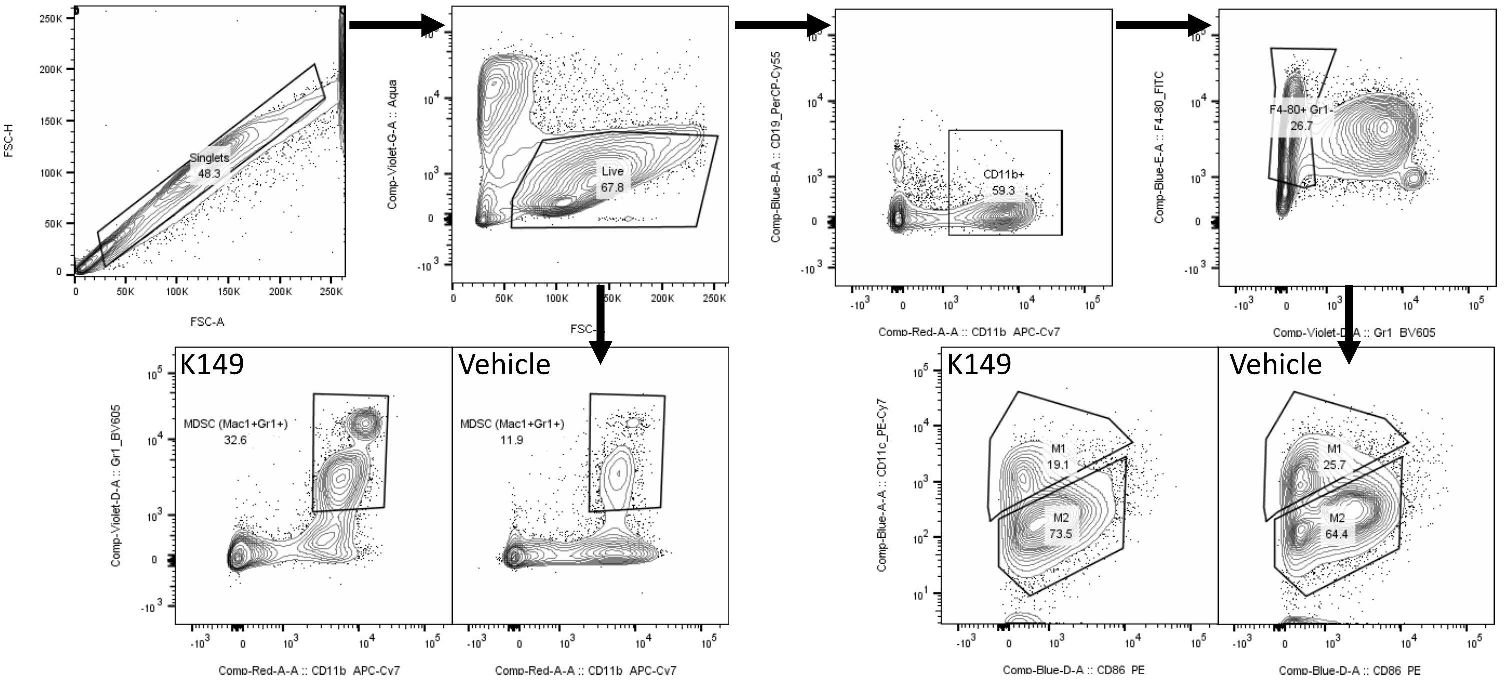


Supplemental Fig.1B. Gating strategy MDSC and M1/M2 K149 Prevention Model

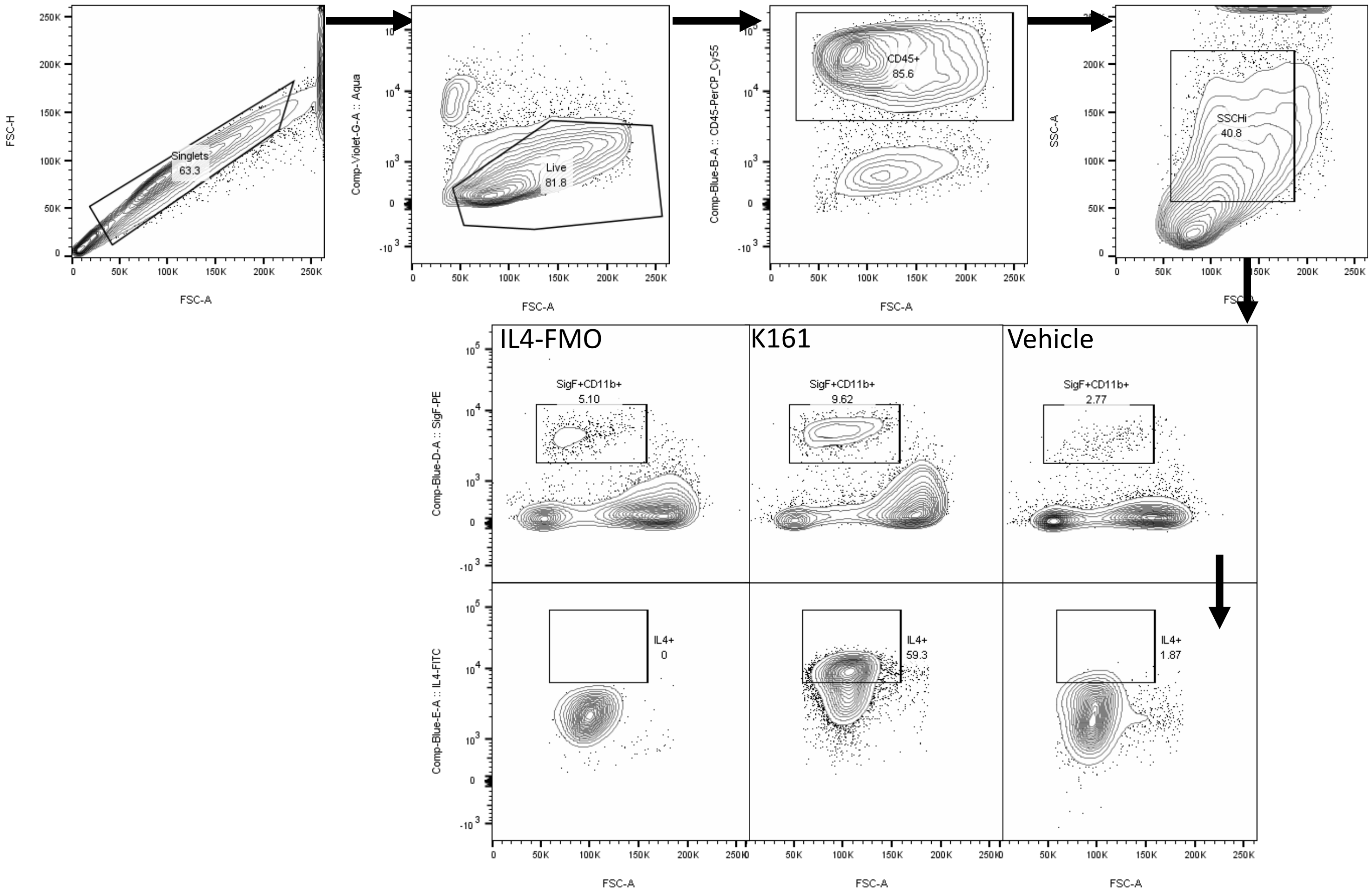


Supplemental Fig.1C Gating strategy Eosinophils K161 Treatment Model

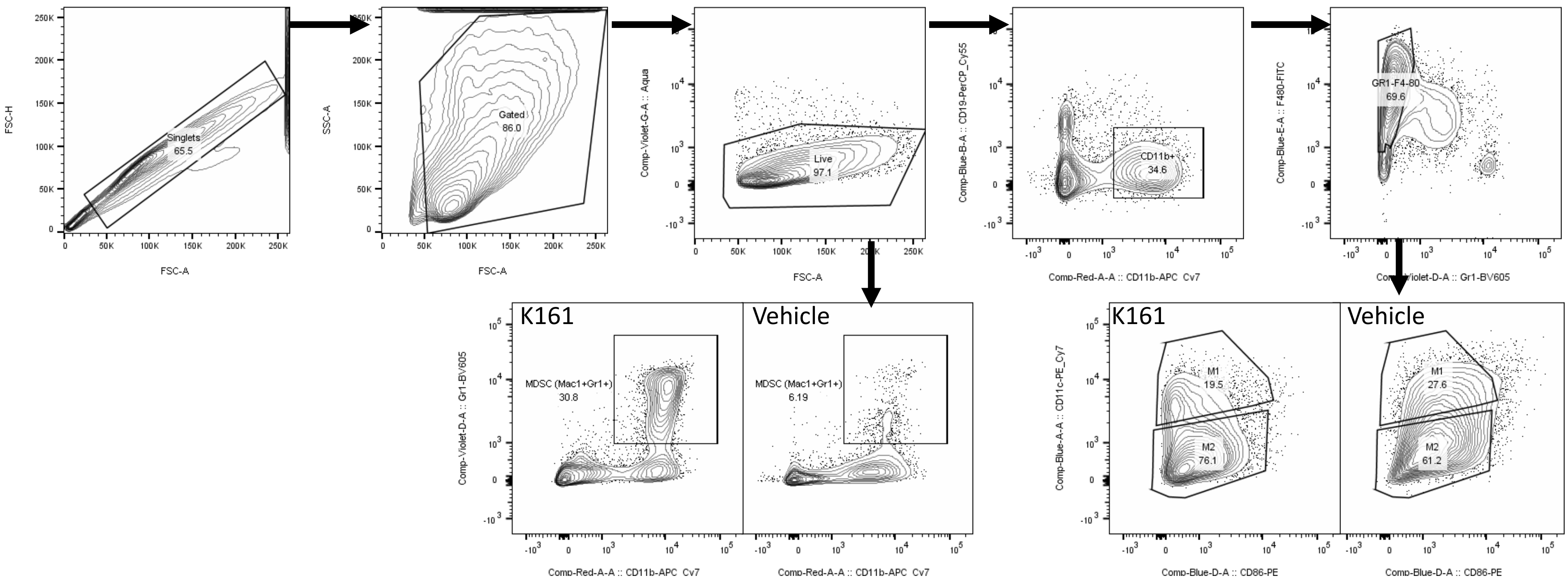


Supplemental Fig.1D. Gating strategy MDSC and M1/M2 K161 Treatment Model
